# A network of steroid receptor transcription factors regulates ovarian chromatin remodelling in the transition to ovulation

**DOI:** 10.64898/2025.11.30.691444

**Authors:** Doan T Dinh, Rebecca L Robker, Darryl L Russell

## Abstract

Steroid receptors are transcription factors activated by progesterone (PGR), androgen (AR) and glucocorticoid (GR), with shared canonical DNA binding sequence. In the ovary, PGR is the key determinant of ovulation while AR and GR play important roles in growing follicles. However, the mechanism that defines the unique physiological roles of these conserved receptors, and whether their functions overlap remains elusive. We investigated the relationship between AR, GR and PGR during folliculogenesis and ovulation. In response to ovulatory hormones, PGR and GR jointly gained binding to novel chromatin sites and had a substantial effect on periovulatory gene regulation, whereas AR-chromatin interactions were repressed. Two modes of PGR action to drive gene activation were identified. Induction of PGR leads to cooperative PGR/GR recruitment to novel promoters, increased histone acetylation and chromatin accessibility, culminating in transcription activation, with PGR being the key component in the unique ovulatory transcriptional complex. Alternatively, PGR tethered to enhancers interacts with pre-accessible, AR/GR-bound promoters to promote gene activation. Our findings illustrate the multi-faceted ovarian steroid receptor interactions which explain how the progressive change in steroid environments throughout folliculogenesis programs granulosa cells during the transition to ovulation.

## Introduction

The NR3C nuclear receptors are steroid-responsive, DNA-binding transcription factors (TF) including progesterone receptor (PGR), androgen receptor (AR), glucocorticoid receptor (GR) and mineralocorticoid receptor (MR), all of which have highly homologous structure. In particular, the DNA binding domains of this family share >85% amino acid sequence identity and recognise a common consensus DNA response element (NR3C motif – GnACAnnnTGTnC). Their distinct functions are thought to be mediated through cell-specific expression of these receptors; however, in granulosa cells of ovarian follicles all are concurrently expressed. Strong evidence indicates PGR, AR and GR and their ligands are each critically involved in maturation of ovarian follicles as well as rupture and ovulation of oocytes (1–4) but their specific molecular actions in mediating this physiological process is unknown. Steroid hormones are important in both anovulatory disorders (5) and contraception (progestins), thus understanding the molecular control of ovulation and female fertility is needed, and for discovery of therapeutic targets for infertility treatments as well as ovulation-blocking contraceptives.

The general transcriptional regulatory mechanisms of steroid receptors are well known to require several important steps: ligand binding releases receptors from the heat shock complex, dimerization and binding to specific chromatin sites, either through the NR3C motif or through tethering with other transcription factors. Interaction with co-regulators promotes the assembly of a transcriptional complex with chromatin remodelling and transcriptional initiation capabilities. The repertoire of expressed co-regulators can establish responses specific to tissue contexts, and recent evidence has indicated that steroid receptors can engage in extensive crosstalk when co-expressed. For instance, in breast cancer, PGR and GR bind mutual chromatin sites, in which GR has a suppressive effect on PGR-driven oncogene transcription (6). In prostate cancer, GR can replace AR at target chromatin sites, which contributes to therapy resistance and poorer prognosis (7). Alternatively, abnormally elevated AR also interferes with GR action by recruiting GR to non-canonical binding sites (8). A tethering mechanism between GR and MR has also been described in multiple contexts (9,10). Further, steroid receptor transcriptional regulatory actions often work through enhancer interaction with distal genes (11). The organisation of chromatin into cell-specific conformational structures creates topological associations of genes and unique regulatory regions that plays a key role in controlling gene expression in different activation states of cells (12). One crucial missing link in hormonal control of ovarian function is understanding the impact on chromatin-chromatin interactions that enables gene regulation through engaging promoters and distal genomic enhancers. While the role of enhancers on transcription regulation in granulosa cells has been described (13,14); the complete enhancer profile in peri-ovulatory granulosa cells and the interaction between enhancers and promoters remains unknown.

Ovulation is a critical differentiation event in ovarian follicles that is triggered by the luteinizing hormone (LH) surge promoting global changes in chromatin accessibility (2), histone modification (15) and different waves of transcription factor activation (16). In granulosa cells, PGR, AR and GR are present during the peri-ovulatory window and their normal functions are important for follicles to progress from folliculogenesis to ovulation. The loss of PGR results in complete anovulation and infertility in female mice (17,18). Antagonists of PGR action suppress ovulation in rodents, macaques and humans (19–22). GR also participates in ovulation mediation, and while it is unknown whether GR is obligatory for ovulation, it has been suggested that cortisol / cortisone ratio is important to follicle survival and oocyte quality in primates (3), and prolonged GR agonist treatment leads to loss of the responsiveness to LH in granulosa cells, thereby affecting ovulatory gene expression (23). AR plays a prominent role in early folliculogenesis and steroidogenesis (24,25), as well as ensuring the responsiveness to the LH-surge in granulosa cells (26,27). Elevated androgen can induce polycystic ovary syndrome (PCOS) in animal models, and is a clinical diagnostic indicator of PCOS in women which results in the accumulation of immature cystic follicles, ovulation failure and sub-fertility (5). The synergy between these receptors remains unknown and likely has important implications in hormone-mediated ovarian transcription regulation and thus female fertility. Considering that each of these receptor transcription factors binds near identical DNA motifs, it remains unclear, if or how they mediate unique events in folliculogenesis and ovulation. Indeed, the molecular mechanisms of AR, GR and MR in the ovary has received relatively little investigation to date.

To elucidate the relationship between different NR3C steroid receptors in the ovarian context, we determined the expression pattern of each steroid receptor and generated comprehensive AR and GR cistrome datasets from granulosa cells in response to *in vivo* LH stimulus and in PGR-null cells through ChIP-seq. We also identified changes in the chromatin structure in response to LH through Micro-C, and then integrated ChIP-seq, ATAC-seq, Micro-C and RNA-seq datasets to provide the comprehensive portrait of steroid receptor action across the genome that is vital for ovulation. Overall, this is the first extensive characterisation of steroid receptors in the physiologically normal reproductive context, which defines the unique mechanisms of steroid interactions controlling ovarian function and reveals novel mechanisms of steroid receptor crosstalk and regulation of chromatin conformation relevant to endocrine-responsive tissues.

## Methods

### Reagents and antibodies

Unless otherwise stated, reagents were purchased from Sigma-Aldrich (St. Louis, MO, USA).

### Animals and granulosa cell collection

All experiments were approved by The University of Adelaide Animal Ethics Committee and were conducted in accordance with the Australian Code of Practice for the Care and Use of Animals for Scientific Purposes (M-2021-093, M-2023-074). CBA x C57BL/6 F1 (CBAF1) mice were obtained from the Laboratory Animal Services (University of Adelaide). PGR knockout (PGRKO) mutant mice (Pgr^tm1Bwo^) (17) were obtained from the Jackson Laboratory (Bar Harbor, USA) and maintained as an in-house breeding colony. Pgr^tm1Bwo^ mice were genotyped from ear or tail biopsies before allocation to experiments and confirmed from replicate biopsies at the time of tissue collection. Experiments used littermate females of each genotype. All mice were maintained in 12 h light / 12 h dark conditions and given water and rodent chow *ad libitum*.

Female mice at 21-days old were either unstimulated or hormonally stimulated by intraperitoneal injection with eCG (Medix Biochemica, Espoo, Finland) followed by hCG (Pregnyl, Merck Sharp & Dohme, Rahway, USA) at 46h post-eCG. Mice were humanely killed by cervical dislocation and ovaries collected at the indicated time points. Granulosa cells were isolated by repeated puncturing of dissected ovaries with a 26G needle in aMEM media.

### Quantitative real-time PCR

Granulosa cells from three mice were pooled together for RNA extraction. A total of three independent experimental replicates were conducted. RNA was extracted from granulosa cells using RNeasy Mini kit (Qiagen, Chadstone, VIC, Australia) for time course experiment or using Trizol RNA extraction method, including DNase treatment, for PGRKO experiment. cDNA was synthesised from 500 ng extracted RNA using SuperScriptIII Reverse Transcriptase kit (Thermo Fisher) and used for qPCR with Taqman methodology (*Pgr* Taqman assay # Mm00435628_m1, *Ar* # Mm00442688_m1, *Nr3c1* # Mm00433832_m1, *Nr3c2* # Mm01241596_m1). Gene expression in each biological replicate (3 biological replicates per time point) was normalised to *Rpl19* (Mm02601633_g1) and fold change was presented as relative to *Rpl19* level using the dC_T_ method.

### Western blot

Granulosa cells were collected as described above. Protein was extracted using RIPA buffer (Merck), and protein lysate was incubated with benzonase to digest DNA. Protein lysate was denatured in LDS buffer (Thermo Fisher) containing 1µl β-mercaptoethanol and heated to 95°C for 10 minutes. Equal volume of samples was loaded and protein separation was achieved by gel electrophoresis at 165 V for 45 minutes. Protein was transferred onto PVDF membrane and blocked in 5% skim milk. Primary antibodies (AR – ab108341, Abcam, GR – 3660S, Cell Signalling Technology, b-actin – A1978, Merck) were used at 1:1000 dilution and secondary antibodies (HRP – 31460, Thermo Fisher, fluorescent – 925-68073, LI-COR) were used at 1:10,000 dilution. Fluorescent blotting was imaged using Odyssey Imager (LI-COR) and chemiluminescent blotting was imaged using SignalBright Max substrate (Proteintech, Rosemont, USA) and the ChemiDoc MP system (Bio-Rad, Hercules, USA).

### Micro-C

Granulosa cells from CBAF1 female mice were stimulated as above and collected at 0h or 6h post-hCG stimulation. Cells from three mice were pooled per biological replicates, 2 biological replicates per timepoint (total 4 samples). Micro-C libraries were generated using the Dovetail Micro-C kit (Cantata Bio, Scotts Valley, USA), with 2-3 technical replicate libraries generated per sample. Shallow sequencing (30M reads/library) was performed to validate quality and complexity of the libraries. All libraries were deep sequenced at minimum 300M reads/library (∼1B reads/sample). Data analysis followed the Dovetail Genomics Micro-C pipeline. Briefly, sequence quality was assessed using FastQC (28) and sequences were aligned to the mm10 mouse genome using BWA-MEM (29) with the following parameters “-5SP-T0”. Valid ligation events were identified using pairtools (30) with the following parameters “parse --min-mapq 40 --walks-policy 5unique --max-inter-align-gap 30”. Read pairs from technical and biological replicates of the same group (pre-or post-LH) were merged, then sorted and deduplicated. Multi-dimensional contact matrices were generated using Juicertools (31) and visualised using Juicebox (32) or the UCSC Genome Browser (33). For genomic organisation identification, the genome was binned at different resolutions (between 5-50 kb bins). Chromatin loops were called using HICCUPS (34) at 5 kb, 10kb and 25kb resolution by default, and differential loops were identified using HICCUPSdiff. Topologically associated domains (TAD) were identified using Arrowhead (34) at 5kb, 10kb and 50kb resolution using default parameters. A/B compartments were identified using CALDER (35) at 25kb resolution (bin resolution automatically optimised) using default parameters.

### Enhancer identification

To classify chromatin interactions, both anchors of each Micro-C chromatin loop identified as above were overlapped with the transcription start site (TSS) genomic coordinates obtained for the mm10 mouse genome through biomaRt (36), using ChIPpeakAnno (37) with maxgap = 0. Anchors containing at least one TSS were classified as ‘promoter’ end, and anchors containing no TSS were classified as ‘enhancer’ end. ATAC-seq of mouse granulosa cells at 0h and 6h post-LH were previously described (2). ATAC peaks were overlapped with ‘enhancer’ end anchors using ChIPpeakAnno, and putative enhancers were defined as ATAC peaks that overlapped with at least one ‘enhancer’ end anchor. An anchor could host multiple enhancers and an enhancer could be found within multiple anchors and thus participated in multiple chromatin looping events. Motif enrichment analysis was performed using HOMER (38). To identify TF binding at enhancers, TF ChIP-seq peaks were obtained from the ReMap 2022 database (39) for mm10 in all available tissues. For each TF, peaks across all tissues were merged using BEDTools (40) and non-redundant peaks were used as consensus TF binding sites. TF binding sites were overlapped with enhancers using ChIPpeakAnno, the percentage of enhancers with TF occupancy was calculated, from which z-score was computed for hierarchical clustering.

### ChIP-seq

For AR ChIP-seq, granulosa cells from PGRKO female mice were stimulated for superovulation as above and collected at 0h post-hCG (wildtype – WT) and 6h post-hCG stimulation (WT and KO). Three to four biological replicates were obtained for each treatment/genotype, each replicate consisted of minimum 1×10^7^ cells pooled from 5 mice. For GR ChIP-seq, granulosa cells from CBAF1 female mice were stimulated as above and collected at 0h or 6h post-hCG, and cells from PGRKO or WT female mice collected at 6h post-hCG, with two biological replicates obtained for each treatment. H3K27ac ChIP-seq was performed on granulosa cells from 44h post-eCG CBAF1 female mice as described above (1 biological replicate). ChIP-seq was performed by Active Motif (Carlsbad, USA) as previously described using AR antibody (#39781, Active Motif), GR antibody (sc-8992, Santa Cruz Biotechnology) or H3K27ac antibody (#39133, Active Motif).

For all datasets, sequences were aligned to the mm10 mouse genome using Bowtie2 algorithm (41). Peak calling from read count followed the algorithm for MACS2 (42) with a p-value cut-off = 10^-10^. Differential binding analysis was performed using DiffBind (43) with DESeq2 linear model, with differentially enriched sites determined to have FDR ≤ 0.05 (LH-induced with logFC > 0. LH-repressed with logFC < 0). Genomic distribution was performed using the ChIPseeker package (44). Motif enrichment analysis was performed using HOMER (38). Visualisation of ChIP-seq signal was through the UCSC Genome Browser. Gene Ontology enrichment analysis was performed using clusterProfiler package (45). Signal heatmap was generated through deepTools2 (46).

### Additional datasets

Additional sequencing datasets were previously generated or obtained from Gene Expression Omnibus (GEO) (Supplementary Table 1). For microarray datasets (oviduct, mammary), differential expression was performed using GEO2R tool (NCBI) at default parameters. For RNA-seq dataset (uterus), differential analysis results from the original publication were used.

## Statistical analysis

Statistical analysis was performed using GraphPad Prism 9 or R. For comparison of qPCR results, one-way ANOVA was used as indicated in the figure legends and statistical significance was considered as p-value ≤ 0.05. For comparison of RNA-seq / ATAC-seq / ChIP-seq difference (differential logFC) between genomic subsets, Wilcoxon signed-rank test with Benjamini-Hochberg correction was performed due to unequal Gaussian residual distribution between groups. Statistically significant comparison is considered to have p-value ≤ 0.001 and |mean logFC| ≥ 0.585 (or 1.5-fold change). Correlation between transcription regulation and TF binding was determined through Pearson correlation.

## Results

### Changes in chromatin interaction in response to ovulation stimulation

To determine the changing chromatin-chromatin interactions in response to the ovulatory LH stimulus, we performed Micro-C on granulosa cells that were pre-(0h) or peri-ovulatory (6h post-hCG). Chromatin-chromatin interactions detected in granulosa cells were largely intra-chromosomal (SFig 1A-B). To determine whether the 3D chromatin landscape corresponds to other genomic changes in peri-ovulatory granulosa cells, including chromatin opening, histone modification and gene expression, we performed compartmentalisation analysis on Micro-C data to identify A and B nuclear compartments, which correspond respectively to active and inactive chromatin regions. Genomic bins were grouped into four A/B states according to change in response to LH-stimulation – A>A (active across ovulation), B>B (inactive across ovulation), B>A (active post-LH only) and A>B (inactive post-LH only). While much of the genome maintained a constant A or B status across ovulation (A>A or B>B), 7% of the genome switched between active and inactive state in response to the LH-stimulus, with similar proportions switching to active (B>A state) or inactive (A>B state) (SFig 1C).

**Figure 1:**
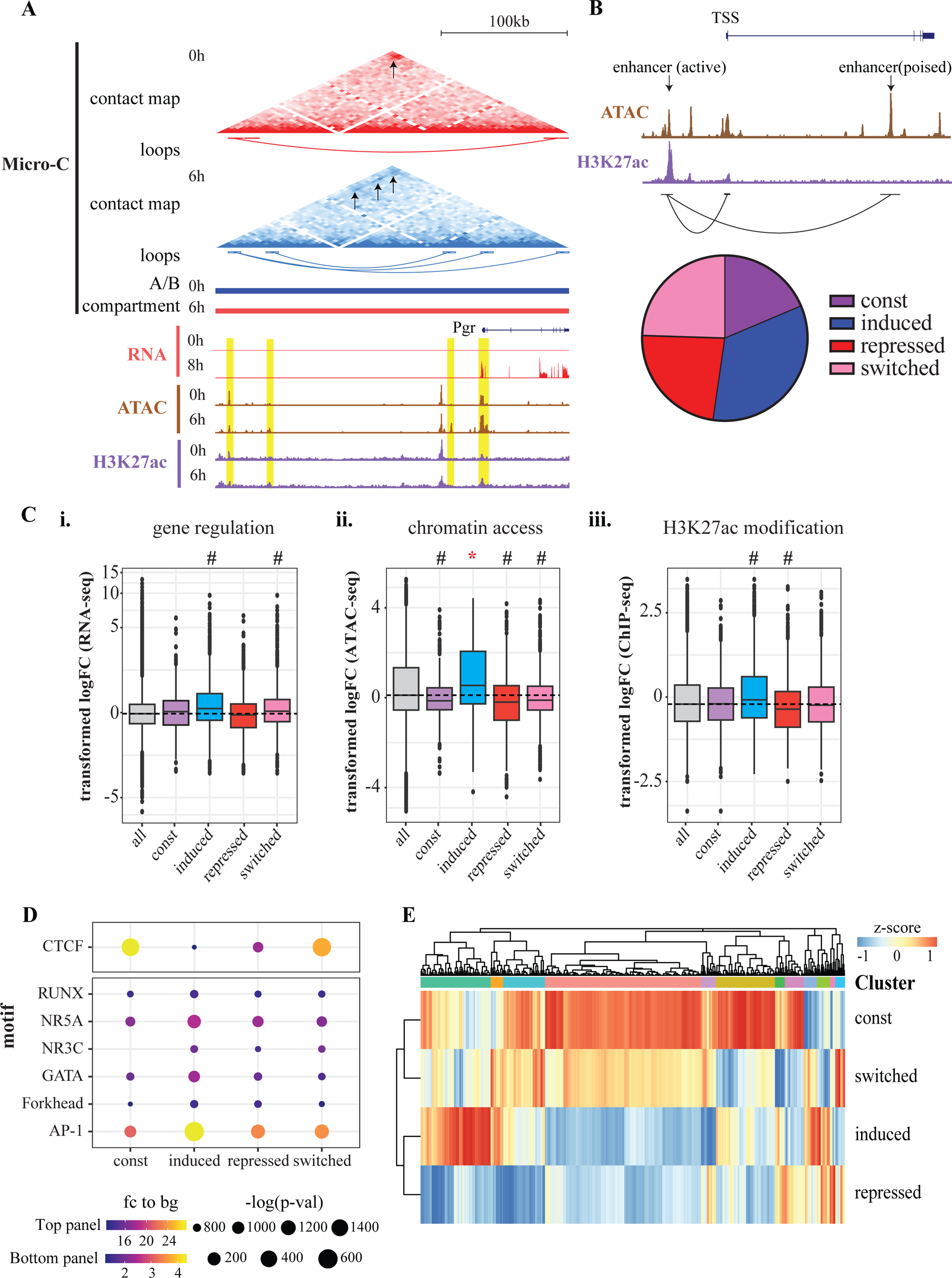
C**h**anges **in chromatin interaction in response to LH-stimulus** (A) Chromatin interaction patterns shown through Micro-C in mouse granulosa cells at 0h vs 6h post-LH stimulus. View window is the topologically associated domain containing the *Pgr* gene locus. From top: chromatin-chromatin contact matrix heatmap, chromatin loops and A/B compartment, genome browser graphical tracks showing gene expression level via RNA-seq, chromatin accessibility via ATAC-seq, H3K27ac occupancy via ChIP-seq. Heatmap: arrows indicate enriched contact points between chromatin sites. A/B compartments: red indicates A (active) compartment, blue indicates B (inactive) compartment. Yellow highlights indicate chromatin loop ends associated with transcriptionally active sites. (B) Representation of enhancer definitions (top) and proportions of identified enhancers in const, induced, repressed and switched groups (bottom) (C) Changes in LH-regulated transcription of genes associated with enhancers (i), chromatin accessibility (ii) or H3K27ac signal (iii) at enhancers in each group (const, induced, repressed, switched). Y axis shows logFC calculated from 8h vs 0h post-LH stimulus for RNA-seq (i), 6h vs 0h for ATAC-seq (ii) and H3K27ac ChIP-seq (iii). Dashed line indicates the median of all identified genes or genomic peaks (‘all’ column). Statistical comparison shown for each enhancer group vs ‘all’, pairwise Wilcoxon test. #: p-value ≤ 0.001, *: p-value ≤ 0.001 and |logFC| ≥ 0.585. (D) Most enriched known transcription factor motifs in each enhancer group. Dot colour indicates fold enrichment to background sequences. Dot size indicates-log(p-value). (E) Heatmap of TF binding preference across four enhancer groups. Z-score is calculated from % of enhancers with each TF binding and clustered. For details on top TF in each clusters, see SFig 1F.

We took advantage of our granulosa cell RNA-seq, ATAC-seq and H3K27ac ChIP-seq data from mice with or without LH-stimulus (2) to determine the net impact of change in A/B state on genomic remodelling and transcription, and defined significantly different groups as those with p-value ≤ 0.001 and |mean logFC| ≥ 0.585 (or 1.5-fold difference) compared to median level. Genomic regions becoming active post-LH showed marked increase in LH-dependent transcription (based on RNA-seq (2), mean logFC = 0.7) and chromatin accessibility (based on ATAC-seq (2), mean logFC = 0.67), and moderate H3K27ac modification (based on ChIP-seq (47), mean logFC = 0.35) (SFig 1D), whereas switching to inactive genomic regions post-LH had the opposite trends. This indicates that changing chromatin interactions is a strong mechanistic determinant of transcriptional regulation. Therefore, Micro-C data gave a robust mechanistic insight on gene regulation in response to LH-stimulus through dynamic contacts between enhancers and TSS. This is demonstrated at the locus of the *Pgr* gene, the key ovulatory determinant that is massively induced post-LH in granulosa cells (Fig 1A). In response to LH-stimulus, the *Pgr* locus showed an overall switch from inactive to active state after LH-stimulus. Additionally, the *Pgr* promoter gained new interactions with upstream chromatin regions (up to ∼200kb away) that were transcriptionally active (accessible and occupied by acetylated histone) (Fig 1A), which might act as distal enhancer to mediate *Pgr* activation.

### Enhancer identification in granulosa cells

Chromatin loop analysis was used to identify interactions between chromatin sites, including between enhancers and their target gene TSS. Among 17,665 significant chromatin loops detected in granulosa cells, approximately one-third was formed in response to LH-stimulus and one-third was lost (SFig 1E), indicating substantial shift in chromatin interactions mediate the dramatic transcriptional reprogramming during ovulation. We integrated information on chromatin access and chromatin-chromatin interaction to identify enhancers in peri-ovulatory granulosa cells. Enhancers were defined as non-TSS chromatin (i.e beyond 3kb to a gene TSS) with high accessibility (significant ATAC-seq peaks) that associate with other enhancers or with gene TSS (Fig 1B). Both active (H3K27ac-positive) and poised (H3K27ac-negative) enhancers were included in this definition. This definition identified a total of 12,660 enhancers, of which one-third (4,262) were found to form chromatin loops only after LH-stimulus (induced enhancers), one-quarter (3,100) switched between pre-and post-LH chromatin loops (switched enhancers), and the remainder were either specific to pre-LH only (repressed enhancers) or remained unchanged (const enhancers).

To determine whether these enhancer categories were associated with different transcriptional regulatory trends, we calculated the net fold change of genes associated with each enhancer group. Genes associated with induced enhancers were more likely to be up-regulated post-LH, compared to global level (mean logFC = 0.39) (Fig 1C). Conversely, genes associated with repressed or constitutive enhancers were not differently expressed compared to global level. Induced enhancers were associated with chromatin reorganisation, with the net fold change in chromatin accessibility being significantly increased for induced enhancers (mean logFC = 0.71) without substantial change in histone acetylation (mean logFC = 0.06). On the other hand, association with repressed or switched enhancers resulted in more blunted or no impact on gene regulation, chromatin accessibility and histone acetylation. Overall, LH-activated *de novo* enhancers contribute more to genomic remodelling and gene activation.

In addition to an enrichment for the binding motif of the classic chromatin organiser CTCF across all enhancer sets, motif analysis identified significant enrichment of the AP-1 TF family motif (including JUN and FOS proteins) at induced enhancers (Fig 1D). Overlaying the binding pattern for mouse TF (determined via ChIP-seq for 648 TF, from the curated ReMap 2022 database (39)) with each enhancer set indicated that const and switched enhancers were occupied by similar sets of TF, whereas induced and repressed enhancers interacted with separate sets of TF (Fig 1E). Further examination of the top enhancer-occupied TF confirmed that CTCF pervasively bound all enhancer subsets, alongside other core components of the transcription complex, including BRD4, CREB1, MED1 and SMARCA4 (SFig 1F-i). TF that preferably bound induced enhancers included members of the AP-1 family (FOS, FOSL2, JUN, JUNB); with the ovulatory determinant PGR also highly enriched (SFig 1F-ii). TF associated with only switched enhancers were weakly enriched overall (<12% enhancers with TF binding) (SFig 1F-iii). These suggest that LH-activated transcription factors (AP-1, PGR) binding to induced enhancers to promote chromatin openness, histone modification and interaction with target genes is an important mechanism for gene activation in response to LH.

### AR and GR chromatin binding dynamics in LH-responsive granulosa cells

Gene and protein expression of the four key NR3C steroid receptors showed dynamic changes in mouse granulosa cells after *in vivo* stimulation of ovulation. *Pgr* transcripts were undetectable in granulosa cells of preovulatory follicles and showed an acute induction within 4 hours post-LH (SFig 2A). *Ar* and *Nr3c1* (GR encoding gene) were abundant in preovulatory granulosa cells with *Ar* showing a downregulation and *Nr3c1* showing modest upregulation after LH-stimulus. *Nr3c2* (MR encoding gene) was low in abundance in granulosa cells at all timepoints. Similar transcriptional expression patterns were observed in response to LH-stimulus in human granulosa cells (SFig 2B). As MR expression was low in granulosa cells, it was not further investigated in this study. The protein levels of AR, GR and PGR corresponded to the mRNA levels (SFig 2C).

ChIP-seq was used to identify granulosa cell AR and GR cistromes before and after LH-stimulus. AR ChIP-seq identified in total 12,030 AR-bound sites, half of which were constitutively bound by AR irrespective of LH-stimulus (Fig 2A). Constitutive AR sites were equally found within proximal promoter regions (within 3 kb of TSS) and non-promoter regions (Fig 2B). Among the AR sites that were LH-regulated, three-quarters were significantly repressed, and the remainder increased after LH-stimulus (Fig 2A). Both LH-induced and LH-repressed sites were highly enriched at intronic and intergenic regions (Fig 2B). To determine the influence of PGR induction on AR binding, AR ChIP-seq was performed comparing PGRWT and PGRKO granulosa cells collected 6h post LH-stimulus. Overall, the absence of PGR had little effect on the granulosa cell AR cistrome (Fig 2A). Changes in AR chromatin binding correspond to changes in the genomic state, and AR binding is highly correlated with chromatin accessibility (Pearson correlation coefficients r = 0.84, p-val < 2.2e-16, Fig 2C) and histone acetylation (r = 0.7, p-val < 2.2e-16, SFig 3A). Motif analysis showed the enrichment of canonical NR3C motif for constitutive and induced AR-bound sites, but especially high enrichment (18-fold over background) in repressed AR sites, indicating that NR3C motif occupancy by AR in preovulatory granulosa cells is repressed by LH-stimulus (Fig 2D). This concurs with our previous observation that the NR3C motif is highly represented in chromatin regions with reduced accessibility post-LH (2). AR in different subsets also showed enriched representation of motifs for AP-1 and NR5A factors, indicating that AR potentially interacts with different sets of co-regulator partners in response to LH-stimulus.

**Figure 2:**
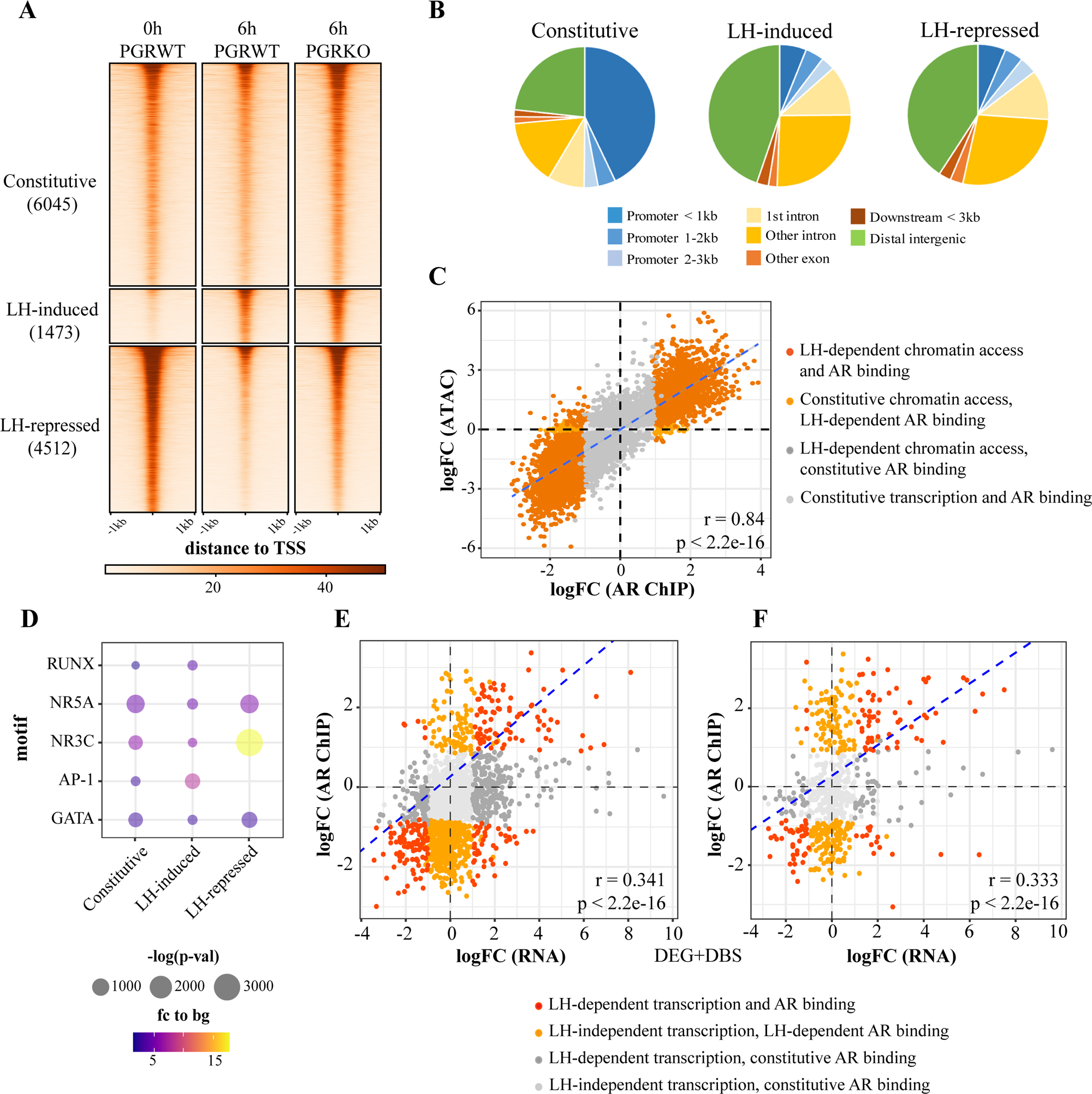
A**R chromatin binding dynamics in LH-responsive granulosa cells** (A) Heatmap of AR ChIP-seq peak signals for granulosa cells from WT 0h or 6h post-LH stimulus and PGRKO after 6h stimulation. AR peaks were partitioned into those that were constitutive, LH-induced or LH-repressed. (B) Genomic distribution of AR peaks in each subset. (C) Correlation between LH-dependent AR ChIP-seq and ATAC-seq signal. Data shown as logFC (ChIP-seq) vs logFC (ATAC-seq). Each dot represents one AR peak, colour labelled for ATAC and/or AR peak classification. Pearson correlation coefficient r = 0.84 (p < 2.2e-16). (D) Top most common known sequence motifs found to be enriched at AR binding sites in each subset. Dot colour indicates fold enrichment to background sequences. Dot size indicates-log(p-value). (E-F) Correlation between LH-dependent gene expression and AR ChIP-seq signal at the promoter of gene (within 3kb of TSS, E) or associated enhancers (F), shown as logFC (RNA-seq) vs logFC (ChIP-seq). Each dot represents one gene, colour labelled for DEG and/or AR binding. Pearson correlation coefficient in promoter r = 0.341 (p < 2.2e-16), in enhancer r = 0.333 (p < 2.2e-16).

We integrated AR ChIP-seq with RNA-seq datasets to investigate the effects on downstream gene expression of AR binding at promoters (within 3kb of the TSS) or at enhancers (defined as above, Fig 1B). AR binding at promoters and enhancers was positively correlated with corresponding gene regulation (r = 0.341 in promoter, r = 0.333 in enhancer, p-value < 2.2e-16) (Fig 2E-F). Gene ontology analysis indicated AR binding at chromatin targets in an LH-dependent manner may regulate different biological processes. Constitutive AR was associated with RNA processing and cell cycle regulation, induced AR was enriched for cell-cell adhesion, immune response and development pathways, and repressed AR was enriched for hormone response and apoptosis (SFig 3B).

GR ChIP-seq identified overall 12,478 binding sites, 80% of which (10,031 sites) were constitutively bound by GR (Fig 3A). Constitutive GR binding showed a strong preference for proximal promoter regions, with more than three-quarters of sites within 1kb of the TSS (Fig 3B). Of the 20% (2,447 sites) that were LH-regulated, 88% (2,146 sites) gained GR binding after LH-stimulus while 12% (301 sites) showed LH-repressed GR binding (Fig 3A). LH-induced and-repressed GR sites were similarly distributed across the genome (Fig 3B). Interestingly, GR binding sites that were gained post-LH were completely lost in the absence of PGR (shown by comparing PGRKO vs PGRWT at 6h post-hCG), indicating that LH-activated PGR facilitates GR recruitment to these new chromatin sites (Fig 3A). In contrast, LH-repressed GR sites were not affected by the lack of PGR (Fig 4A). Similar to AR, GR chromatin binding is highly positively correlated with changes in chromatin opening (r = 0.78, p-val < 2.2e-16) (Fig 3C) and transcriptionally active histone modification (r = 0.64, p-val < 2.2e-16) (SFig 4A). Motif enrichment analysis indicates the canonical NR3C motif (>7.5-fold to background), along with AP-1, NR5A and RUNX, were associated with induced GR regions (Fig 3D). Repressed GR regions were enriched for NR3C and ER binding motifs, and constitutive GR sites were enriched for DNA sequences that are commonly associated with core promoter regions, including the CCAAT and GC-rich boxes (Fig 3D).

**Figure 3:**
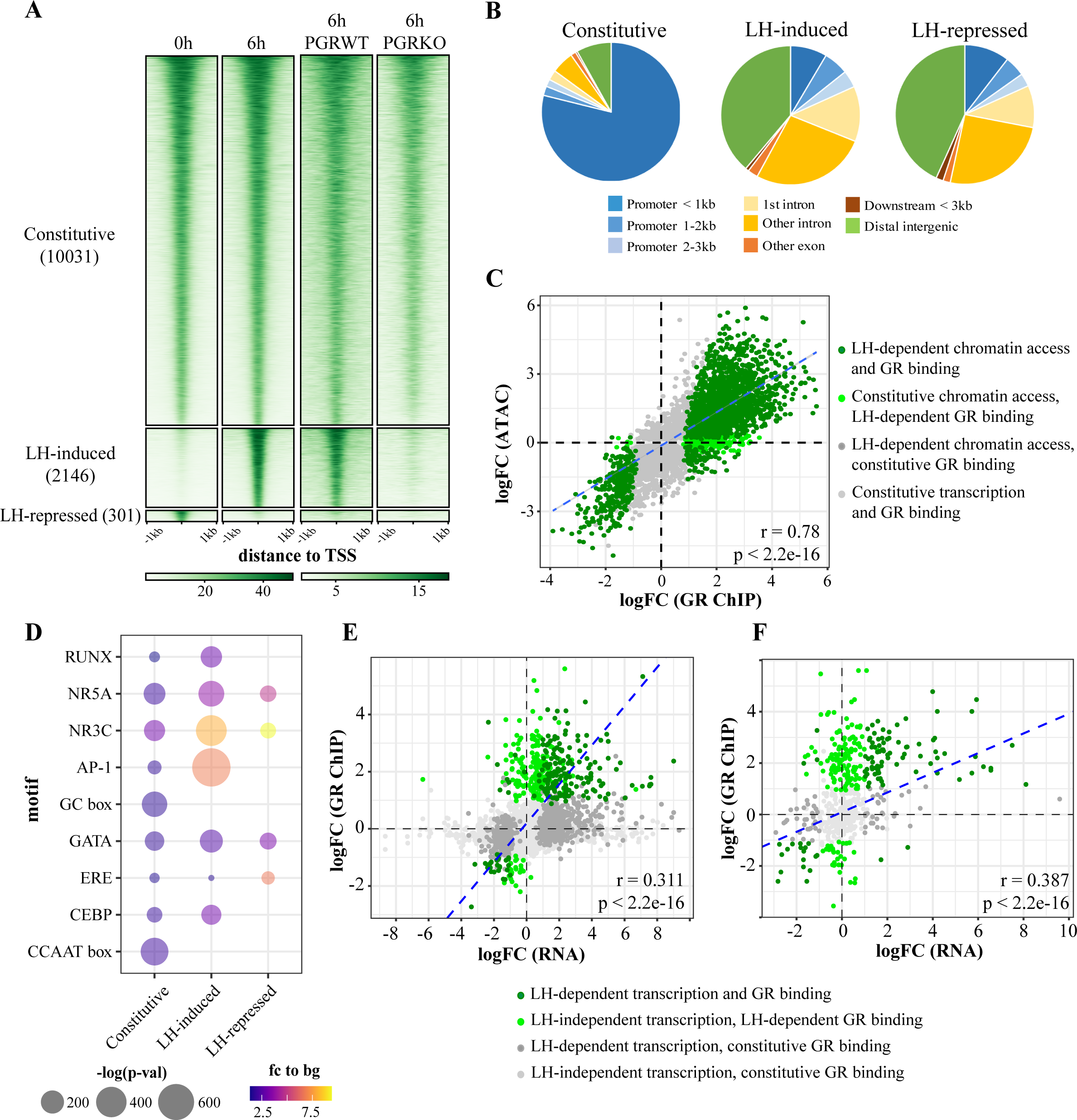
G**R chromatin binding dynamics in LH-responsive granulosa cells** (A) Heatmap of GR ChIP-seq peak signals for granulosa cells 0h or 6h post-LH stimulus and PGRKO after 6h stimulation. GR peaks subset into those that were constitutive, LH-induced or LH-repressed. (B) Genomic distribution of GR peaks in each subset. (C) Correlation between LH-dependent GR ChIP-seq and ATAC-seq signal. Data shown as logFC (ChIP-seq) vs logFC (ATAC-seq). Each dot represents one GR peak, colour labelled for ATAC and/or AR peak classification. Pearson correlation coefficient r = 0.78 (p < 2.2e-16). (D) Top most common known sequence motifs found to be enriched at GR binding sites in each subset. Dot colour indicates fold enrichment to background sequences. Dot size indicates-log(p-value). (E-F) Correlation between LH-dependent gene expression and GR ChIP-seq signal at the promoter of gene (within 3kb of TSS, E) or associated enhancers (F), shown as logFC (RNA-seq) vs logFC (ChIP-seq). Each dot represents one gene, colour labelled for DEG and/or GR binding. Pearson correlation coefficient in promoter r = 0.311 (p < 2.2e-16), in enhancer r = 0.387 (p < 2.2e-16).

**Figure 4:**
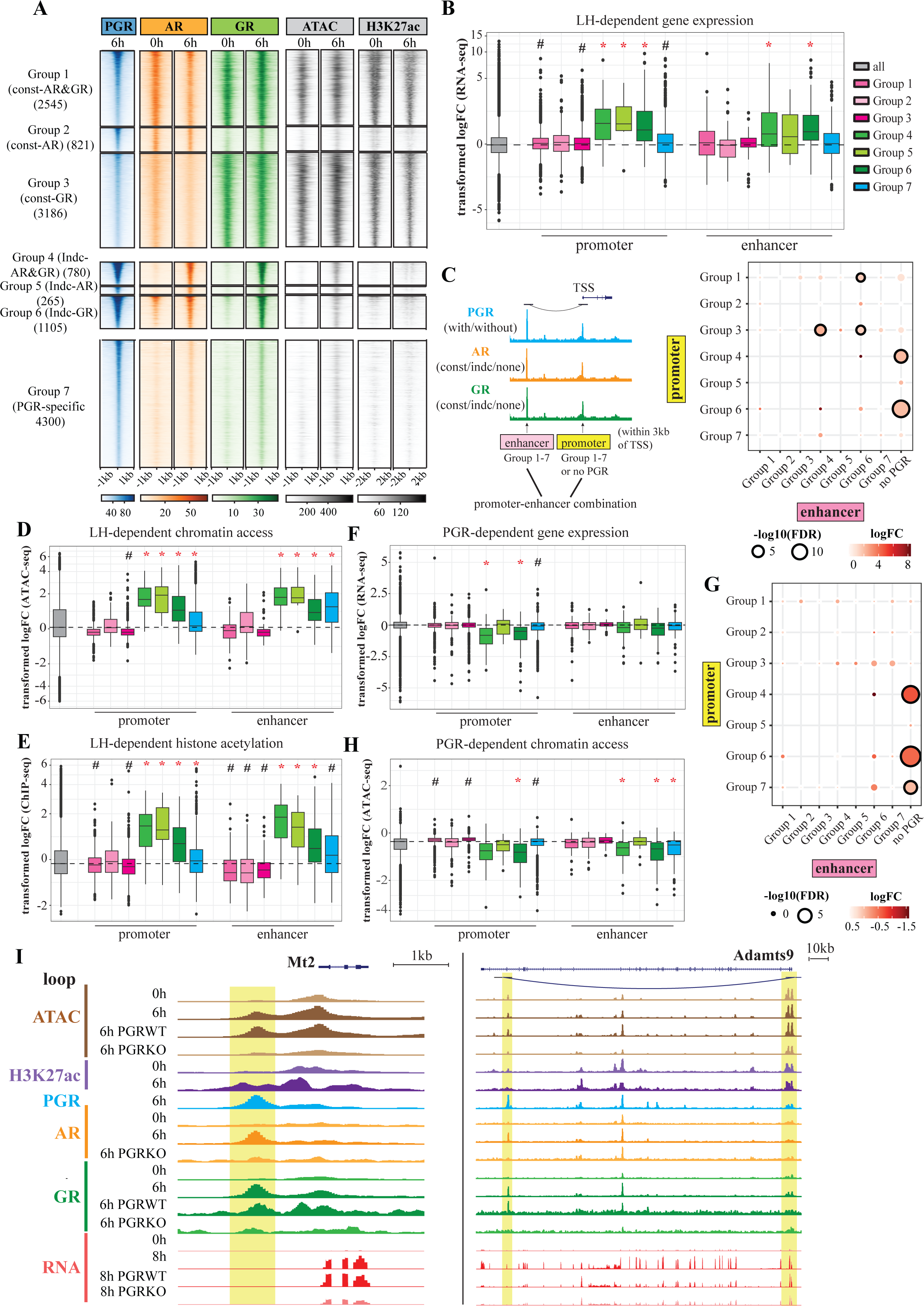
E**f**fect **of steroid receptor co-binding on chromatin and gene regulation** (A) Heatmap of most common subsets of regions with PGR-bound peaks in PGR ChIP-seq, and corresponding ChIP-seq signal for PGR, AR GR, and H3K27ac, and ATAC-seq in 0h/6h post-LH stimulus granulosa cells for each subset. (B) Changes in LH-mediated transcription in each PGR subset. Y axis shows logFC calculated from RNA-seq granulosa cells of 8h vs 0h hCG-treated mice. Data shown for PGR binding in promoters or enhancers. Dashed line indicates the median of all identified genes or genomic peaks (‘all’ column). Statistical comparison shown for each PGR subset vs ‘all’, pairwise Wilcoxon test. #: p-value ≤ 0.001, *: p-value ≤ 0.001 and |logFC| ≥ 0.5855. (C) Definition of promoter/enhancer combination used for synergy analysis, with each promoter or enhancer classified into one of seven PGR subsets (left), and changes in LH-mediated transcription regulation (8h vs 0h post-hCG) in each combination (right). Dot colour indicates mean logFC per combination, dot size indicates-log(FDR) (from pairwise Wilcoxon test). Dots with bold outline are statistically significant (FDR ≤ 0.05). (D-E) Changes in LH-mediated chromatin accessibility (D) and H3K27ac signal (E) in each PGR subset. Y axis shows logFC calculated from 6h vs 0h post-LH ATAC-seq (D) or H3K27ac ChIP-seq (E). Data shown for PGR binding in promoters (i) or enhancers (ii). (F) Changes in PGR-mediated transcription regulation in each PGR subset. Y axis shows logFC calculated from KO vs WT post-LH RNA-seq. Data shown for PGR binding in promoters (i) or enhancers (ii). (G) Synergy analysis of change in PGR-mediated transcription (PGRKO vs PGRWT) in each combination of PGR subset at promoters (y axis) or enhancers (x axis). Dot colour indicates mean logFC per combination, dot size indicates-log(FDR) (from pairwise Wilcoxon test). Dots with bold outline are statistically significant (FDR ≤ 0.05). (H) Changes PGR-mediated chromatin accessibility in each PGR subset. Y axis shows logFC calculated from KO vs WT ATAC-seq. Data shown for PGR binding in promoters (i) or enhancers (ii). (I) Example of genes in which PGR facilitates gene expression through promoter binding (*Mt2*, left) or enhancer (*Adamts9*, right). Yellow highlight indicates PGR binding regions associated with genes.

GR binding was shown to have substantial effect on transcription regulation in response to the LH-surge. Overall, GR binding both at promoters and gene-associated enhancers was positively correlated with corresponding gene regulation (Pearson correlation coefficient r = 0.311 in promoter, r = 0.387 in enhancer, p-value < 2.2e-16) (Fig 3E-F). Each GR bound subset was also associated with different biological functions, as shown through gene ontology analysis. GR bound genes were associated with transcription regulation, immune response, apoptosis, and cell adhesion; and repressed GR was associated with reproductive pathways (SFig 4B).

### LH-induced steroid receptor interaction facilitates chromatin remodelling and gene activation

ChIP-seq for AR (Fig 2A), GR (Fig 3A) and PGR (47) demonstrate that chromatin interactions of all three steroid receptors are in fact regulated by ovulatory stimulus, with GR binding to new sites appearing to be dependent on PGR. Cross-comparison of the cistromes of all three receptors in granulosa cells indicates substantial overlap (SFig 5A). Of the 13,382 PGR peaks, 9,082 were also bound by AR and/or GR, with the majority consisting of constitutive or LH-induced AR and/or GR. Only 380 PGR peaks were also LH-repressed AR/GR, suggesting that PGR does not compete or displace AR and GR on the chromatin, but rather the activation of PGR after LH-stimulus induces AR and GR binding at new sites previously inaccessible to these receptors. We next examined the relationship between steroid receptors in peri-ovulatory granulosa cells and classified PGR binding sites into seven subsets based on the LH-responsive chromatin binding of AR and GR (Fig 4A). PGR binding sites with constitutively bound GR with or without AR (Groups 1, 3) showed strong sustained ATAC and H3K27ac ChIP signals, whereas constitutive AR binding only (Group 2) showed weak ATAC and H3K27ac signal (Fig 4A). LH-induced AR and/or GR at PGR co-bound sites (Groups 4-6; 2,150 sites), were associated with moderate induced ATAC and H3K27ac signals. The remaining one-third of PGR sites, which were not co-bound by AR or GR (Group 7), also showed weak ATAC and H3K27ac signals. Assessing the genomic distribution of PGR-bound subsets showed that sites pre-occupied by GR prior to the LH-stimulus (Groups 1, 3) were overwhelmingly enriched within 1kb of TSS (70-90% of all sites) (SFig 5B). Genes associated with different PGR subsets were also linked to distinct biological functions. (SFig 5C).

Integrating ChIP-seq with RNA-seq and ATAC-seq data revealed that the PGR-GR interaction is most important for ovulatory gene regulation. LH-induced PGR along with AR and/or GR (Groups 4-6) at promoters or enhancers was associated with the most profound LH-mediated gene activation, whereas PGR co-binding at constitutive AR and/or GR (Groups 1-3) or without AR/GR (Group 7) had little impact on gene activation (Fig 4B). The synergy between PGR promoter-and enhancer-binding on gene activation was examined by comparing transcriptional changes between genes with different combination of PGR subsets at their promoter and associated enhancers (Fig 4C). Surprisingly, genes in Group 4 and Group 6, which exhibited high levels of LH-dependent activation (Fig 4B), did not benefit from additional PGR binding at associated enhancers (Fig 4C). On the other hand, genes with PGR and constitutive AR & GR (Group 1), or constitutive GR (Group 3) gained transcription activation when associated with enhancers with PGR and induced-GR (Groups 4, 6). These enhancers that supported gene activation were largely activated post-LH only, rather than constitutive active enhancers (SFig 6A). Both chromatin accessibility and histone acetylation were significantly increased at promoters and enhancers associated with induced AR and/or GR subsets (Group 4-6) (Fig 4D-E). Motif analysis of PGR/indc-GR binding sites (Groups 4, 6) associated with promoters or enhancers indicates differences in the underlying mechanisms in these two groups. PGR binding at promoters was largely associated with the presence of canonical NR3C motifs, which were absent at PGR-bound enhancers (SFig 6B). Rather, motifs for AP-1, GATA, NR5A and RUNX TF families were found at enhancers, suggesting that PGR is recruited to enhancers through tethering rather than direct DNA binding. Surveying LRH-1 (or NR5A2, member of the NR5A family) and RUNX1 (member of the RUNX family), which play important roles in gene regulation in ovulation (2,48) showed that both TF gain chromatin binding at PGR-bound promoters and enhancers in response to the LH-surge (SFig 6C). The data indicate that LH-induced PGR promotes new GR binding at promoters or LH-activated enhancers, and this PGR-GR interaction enables genome remodelling and transcription regulation during ovulation.

Integrating RNA-seq and ATAC-seq from granulosa cells of PGRKO and WT mice illustrated the importance of PGR in these mechanisms. The activation of genes with induced-GR binding in promoters (Groups 4, 6) was reduced in the absence of PGR; however, genes with induced-AR binding (Group 5) were unaffected (Fig 4F). These genes are not PGR-regulated in other PGR-positive female reproductive tissues (oviduct, uterus, mammary tissues) (SFig 6D-F), suggesting that LH-induced PGR/GR interaction is uniquely important for granulosa-specific gene activation. While the loss of PGR binding at promoters significantly affected gene activation in granulosa cells, the absence of PGR did not affect genes regulated through enhancers (Fig 4F). This is further highlighted when the synergy between promoter-enhancer PGR action was examined in PGRKO vs WT, in which only genes with PGR/induced-GR co-binding at promoters were dependent on the presence of PGR for activation (Groups 4, 6) whereas genes in all other subsets, irrespective of PGR occupancy at enhancers, were largely unaltered (Fig 4G). The accessibility of chromatin sites harbouring PGR (Groups 4, 6) at promoters or enhancers was reduced in PGRKO (Fig 4H), indicating PGR is required for chromatin opening. Overall, PGR transcriptional activation in ovulatory granulosa cells is associated with LH-induced GR chromatin binding, with two possible modes of action. Mainly, PGR binding at promoters is essential for the recruitment of GR, genomic remodelling and gene activation (e.g seen in *Mt2*, Fig 4I). Alternatively, PGR can also promote expression of pre-accessible genes through interaction with LH-activated enhancers (e.g seen in *Adamts9*); however, the presence of PGR is dispensable.

## Discussion

AR, GR and PGR play diverse important roles in many biological contexts, including key processes in the ovary leading to ovulation. Such roles are achieved through receptor-specific interactomes, including complex inter-relationships among the receptors themselves, which profoundly affect hormone responses and physiological consequences (49–51). By analyzing steroid receptor cistromes in conjunction with comprehensive characterisation of the granulosa cell genome responses to ovulatory cues *in vivo*, we have elucidated fundamental differences in the underlying mechanisms of each steroid receptor that helps to explain stark contrasts in functions. At the same time, the cooperation between steroid receptors illustrates a more intricate hormonal control of the process of ovulation than previously appreciated.

Our data indicates the complex crosstalk of AR and GR with PGR action in the peri-ovulatory period, thus unveils an interactive mechanism of hormonal control of LH-regulated gene expression. Upon activation by the LH-surge, PGR can bind promoters that are inaccessible prior to LH-stimulation through the canonical NR3C motif. This facilitates GR recruitment to these new sites, enables histone acetylation and chromatin opening, leading to transcription activation. In these activated genes, the presence of PGR is indispensable, and the loss of PGR prevents chromatin reorganisation that is required for transcription; GR binding is also a defining event, indicating that these two steroid networks are integrated. Alternatively, some genes with pre-accessible, AR/GR-preoccupied promoters are activated in response to the LH-surge through new association with PGR-bound enhancers. PGR binding at these enhancers likely involves tethering by other TF, including AP-1 and RUNX1, rather than direct DNA binding. While the presence of PGR at these enhancer sites is required for GR recruitment and chromatin opening, PGR at enhancers is expendable for transcription. Distinct groups of genes are regulated by either mechanism; thus, acting in parallel, the two modes of action allow PGR to modulate specific gene sets in granulosa cells.

Both of the described modes of action feature a key mechanism previously not described in other tissue contexts – PGR recruitment of GR to novel chromatin targets not previously accessible to GR. In response to the LH-surge, GR is able to bind new promoters and enhancers, and this mechanism is almost entirely reliant on co-binding with PGR. This suggests that GR may be tethered to new targets by PGR, but PGR-induced chromatin accessibility is also a possible mechanism. Given that GR is expressed in high levels prior to the LH-surge, it remains unclear how GR gains PGR-dependent novel binding sites in response to LH. A dramatic increase cortisol ligand concentration in granulosa cells in response to the LH-surge (52) likely also changes GR activity. GR co-binding a subset of PGR target sites has also been observed in breast cancer (6,50) in which GR modulates PGR action by enhancing PGR chromatin binding at specific genes while sequestering PGR from other targets. Whether PGR reciprocally impacts GR action in mammary or other tissues is unknown; thus, to our knowledge this is the first direct evidence of PGR mediating GR action in any biological context. LH-induced PGR/GR co-binding is associated with changes in chromatin opening, histone modification and the formation of promoter-enhancer chromatin interactions. Through PGRKO, we show that chromatin opening is dependent on the presence of PGR, whether that involves the direct action of PGR or GR (or other tethered TF) is not clear; however, it seems reasonable to conclude the combined action of this unique transcriptional complex is mechanistically important.

Unlike GR, ovarian AR action is more prominent prior to the LH-surge, corresponding to the known roles of AR in follicle development (25). In response to LH-stimulation, AR shows moderate downregulation in expression and a substantial loss in chromatin binding, corresponding to the downregulation of folliculogenesis genes (eg *Cyp19a1*, *Fshr*). This is similar to the pattern observed for estrogen receptors alpha (ERα) and beta (ERβ) in granulosa cells [Dinh et al 2025 manuscript under review]. Indeed, AR overlapped extensively with ERα and ERβ chromatin binding in granulosa cells, especially at sites present prior to LH-stimulus. While some new AR binding sites are gained post-LH, the majority of which are at shared PGR/GR sites, such binding occurs irrespective of the presence of PGR. Additionally, while LH-induced AR/PGR co-binding (with or without GR) contributes to chromatin remodelling and gene activation in response to the LH-surge, such genomic and transcriptomic changes do not require the presence of PGR, suggesting that AR alone can modulate transcription independently of PGR action. Little research has focused on the physical and functional interactions between AR and PGR; in breast cancer, while AR and PGR are shown to extensively share target chromatin sites (50), the implications on gene regulation remain unexplored. Here, we show that while there is considerable overlap between PGR and AR cistromes, this overlap does not contribute significantly to PGR action. Thus, unlike the close association between PGR and GR, the net loss of AR-chromatin binding may be a physiologically relevant event in the transition from folliculogenesis to ovulation.

In addition to cooperation with GR, PGR action in granulosa cells likely also involves interaction with other TFs, especially at PGR/induced-GR enhancers which lack the NR3C motif. Previously, we have identified RUNX1 to be an important contributor to PGR ovulatory action (2). Here, RUNX motif is enriched in both modes of PGR action, and LH-induced RUNX1 is shown to accompany PGR at most promoters and enhancers. Members of the NR5A family (SF-1 and LRH-1) are expressed in granulosa cells, are involved in folliculogenesis and ovulation, and are important for female fertility (48,53,54). In the ovary, LRH-1 can also act in conjunction with steroid receptors such as ERβ (55). The NR5A motif was found at both PGR-occupied promoters and enhancers, and LRH-1 co-bound promoters or enhancers targeted by PGR. The motif for AP-1 (JUN/FOS) family was also highly enriched in both PGR-bound promoter and enhancer sets. AP-1 members are expressed in granulosa cells and have prominent roles in regulating gene expression that are important for ovulation, including steroidogenesis, prostaglandin synthesis and matrix remodelling (56,57). In the human granulosa cell line KGN, we showed RUNX1, GR, JUND and FOSL2 to be in the PGR protein complex (58), supporting RUNX1 and AP-1 factor participation in the PGR/GR granulosa cell transcription complex.

Our results also highlight another feature of the PGR regulatory mechanism, in which either binding of promoter or enhancers alone is sufficient for LH-induced PGR/GR to facilitate chromatin remodelling and gene activation. We found very few instances of LH-induced PGR/GR co-binding at both the promoter and enhancers of the same gene (5 genes in total), suggesting that synergistic promoter/enhancer effect is rare. However, while the ablation of PGR (in PGRKO mice) blocks the transcription of genes with PGR-bound promoters, genes with PGR-bound enhancers were unaffected. Thus, PGR is obligatory for the activation of gene promoters but not enhancers. The role of PGR in mediating transcription through enhancers has previously been described in the regulation of specific target genes, including *Zbtb16* (59), and *Adamts1* (2), and the association of PGR with global enhancers have also been profiled in mammary (60) and endometrial cells (61). However, the link between promoter and enhancer regulatory mechanisms on PGR-mediated gene expression has not previously been investigated. Transcriptional activation in the absence of PGR at enhancers is possibly rescued by compensatory regulatory mechanisms, either involving the action of alternative TF or the activation/interaction with other non-PGR binding enhancer sites. The impact of PGR on three-dimensional chromatin-chromatin interactions in any biological contexts has yet to be determined. Further characterisation of the PGR-dependent chromatin-chromatin interaction map in granulosa cells will be required to fully identify its role in enhancers.

Overall, two distinct regulatory mechanisms for PGR were identified, in which PGR separately mediates gene regulation through interaction with gene promoters or enhancers, facilitation of GR recruitment to chromatin and genomic remodelling. Both mechanisms act in concert to modulate ovulatory gene activation that occurs in response to the LH-surge. As steroid receptors are crucial for normal physiology as well as the aetiology of multiple diseases, especially breast and prostate cancers, this has important implications in the development of steroid receptor-specific therapeutics.

## Data availability

The datasets generated and/or analysed during the current study will be made available upon publication in a peer-reviewed journal.

## Author Contributions Statement

Conceptualization D.T.D., R.L.R, D.L.R., Formal analysis and interpretation D.T.D., R.L.R., D.L.R., Funding acquisition D.L.R., Investigation D.T.D., Supervision D.L.R., Resources R.L.R., D.L.R., Visualization D.T.D., Writing – original draft D.T.D, Writing – review & editing D.T.D., R.L.R., D.L.R.

## Funding

This work was supported by the National Health and Medical Research Council [GNT1110562 to D.L.R., GNT1117976 to R.L.R.]; the Bill and Melinda Gates Foundation [INV-024199 to D.L.R.]; and the Lloyd Cox Obstetrics & Gynaecology Research Fund [to D.T.D.].

## Conflicts of interest

The authors have nothing to disclose

## Supporting information

SFig 1

SFig 5

SFig 2

SFig 3

SFig 4

SFig 6

---

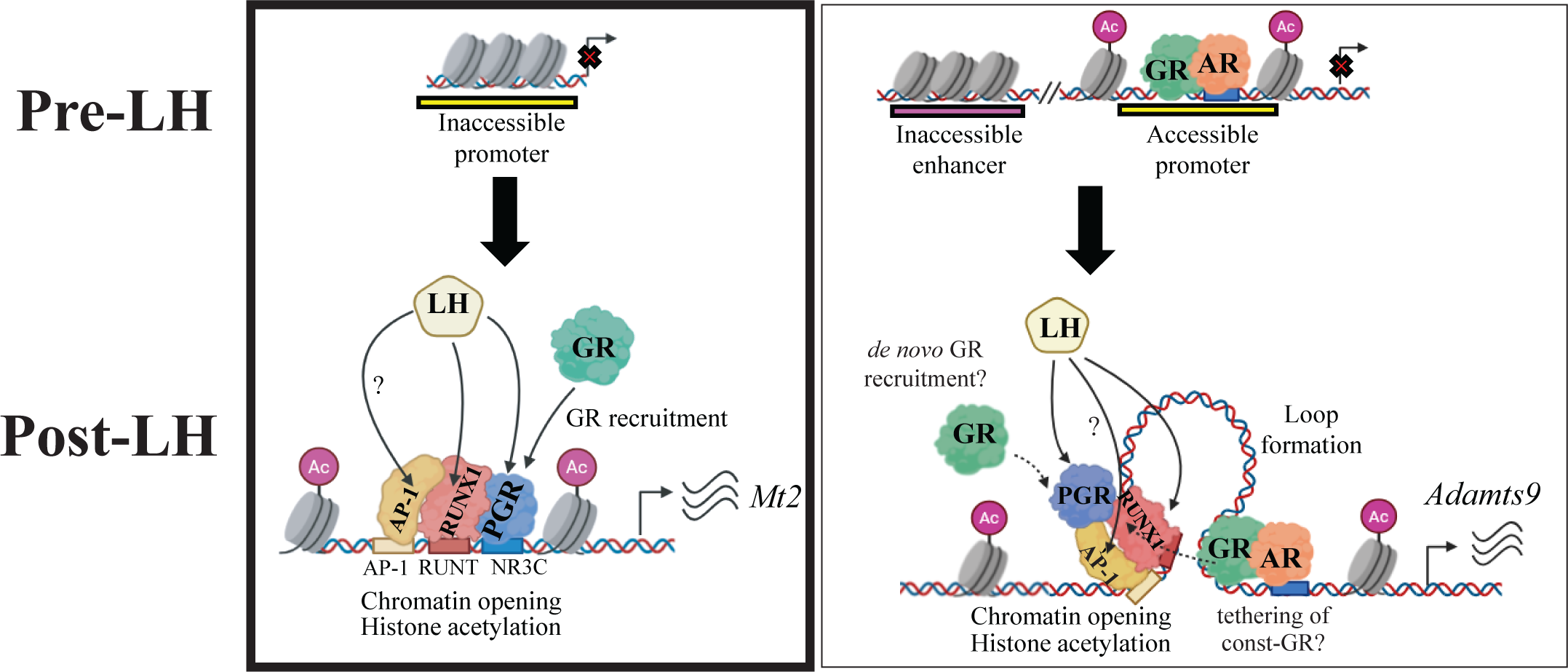

## Notes

### Competing Interest Statement

The authors have declared no competing interest.

