## Supplementary figures and images for "A network of steroid receptor transcription factors regulates ovarian chromatin remodelling in the transition to ovulation"

### SFig 1

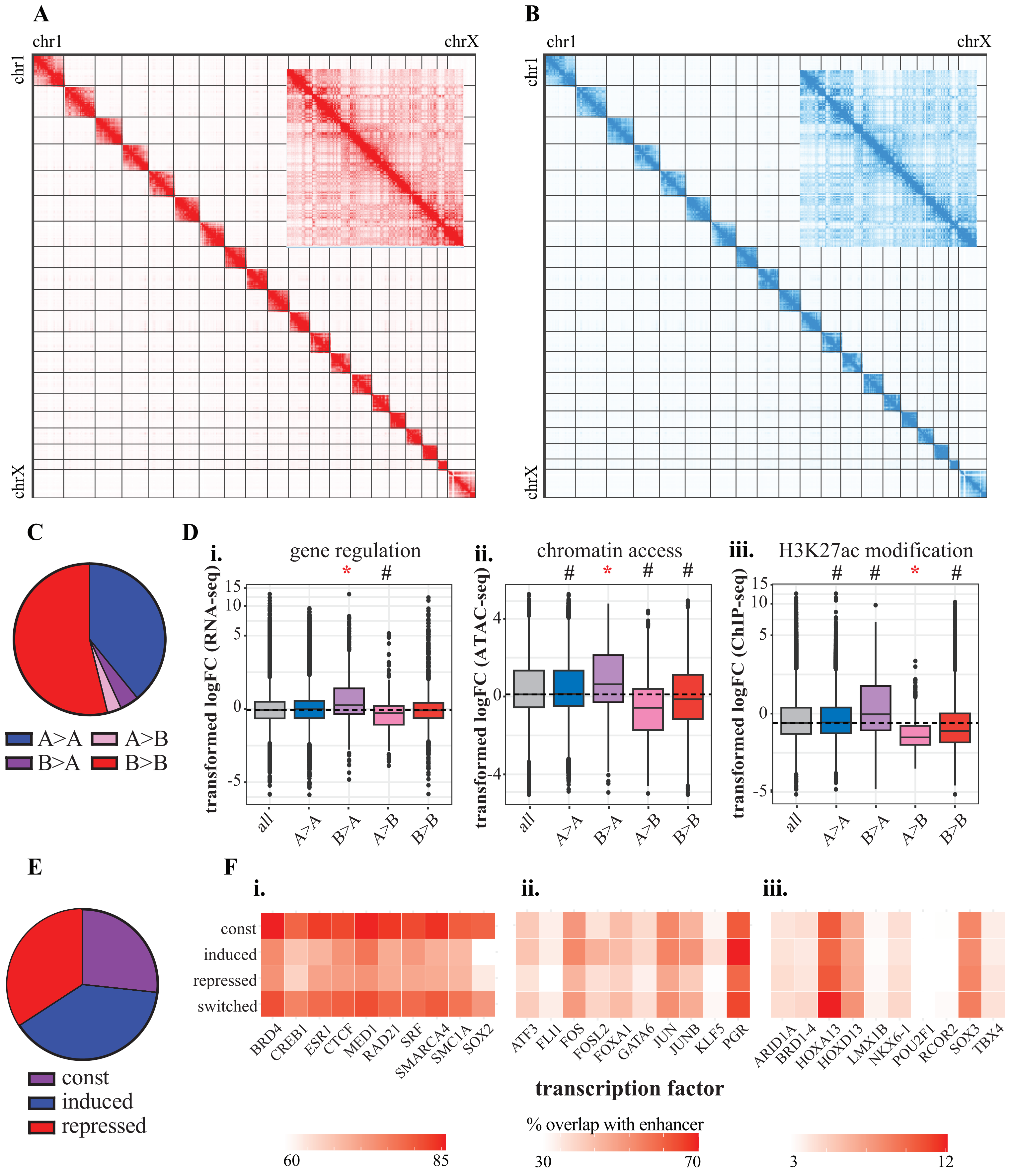

### SFig 2

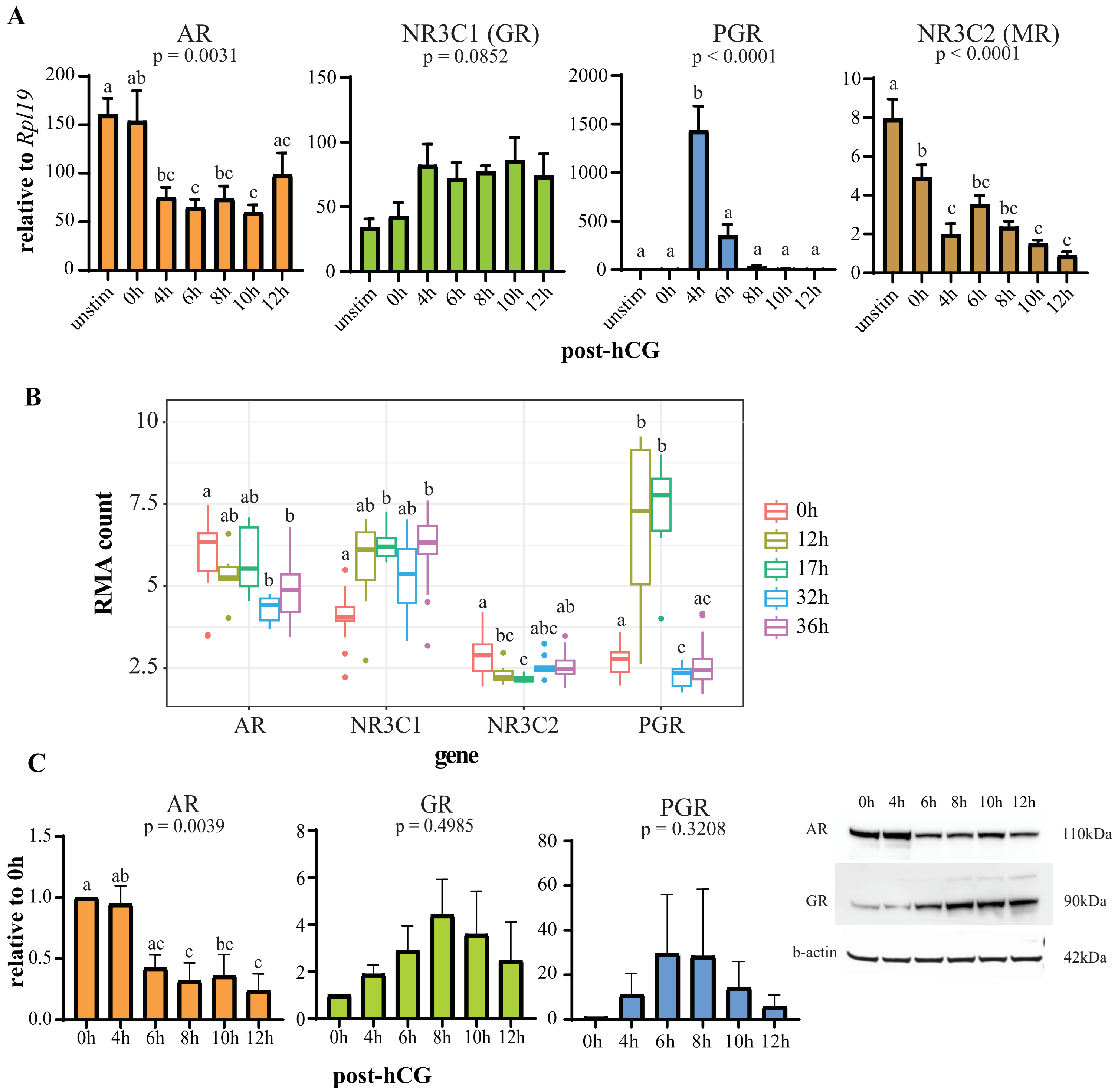

### SFig 3

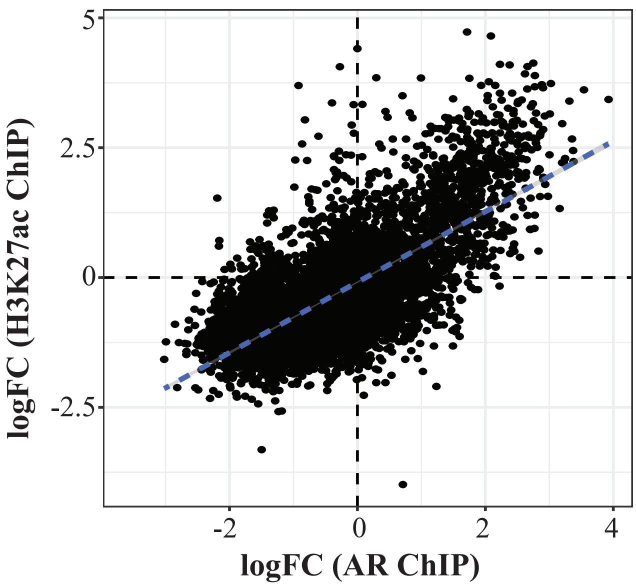

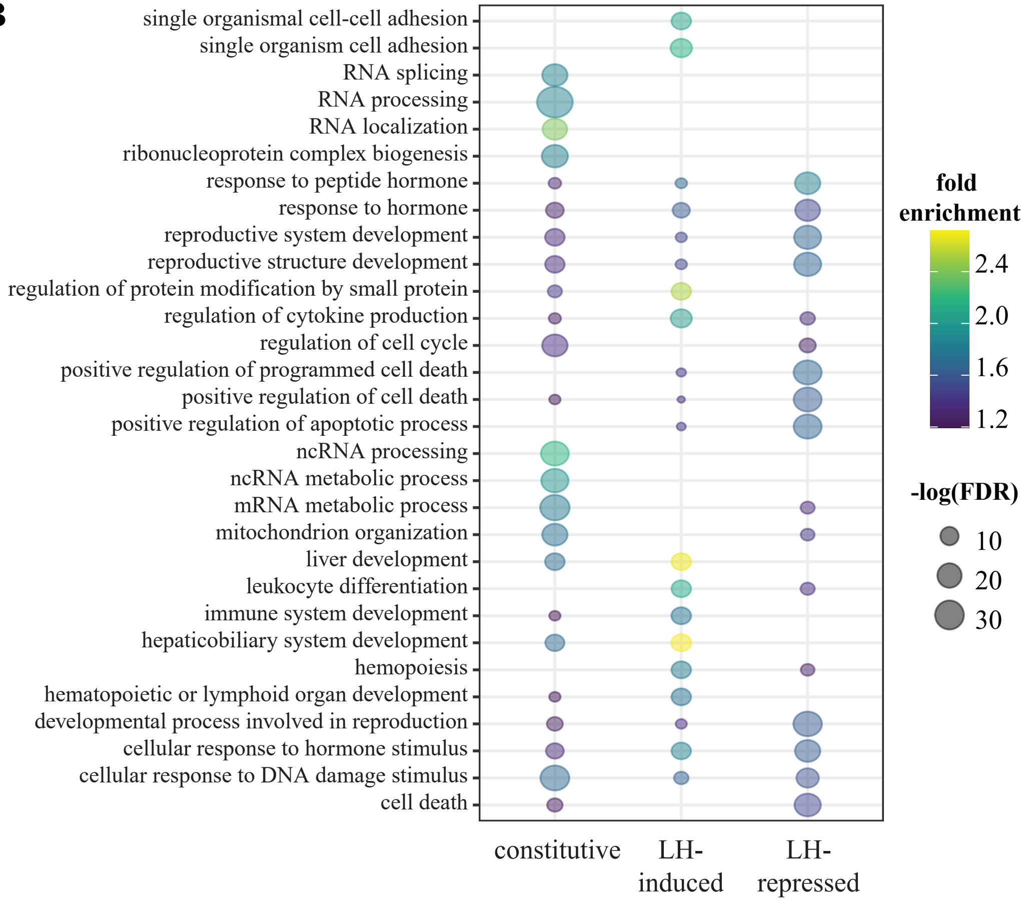

### SFig 4

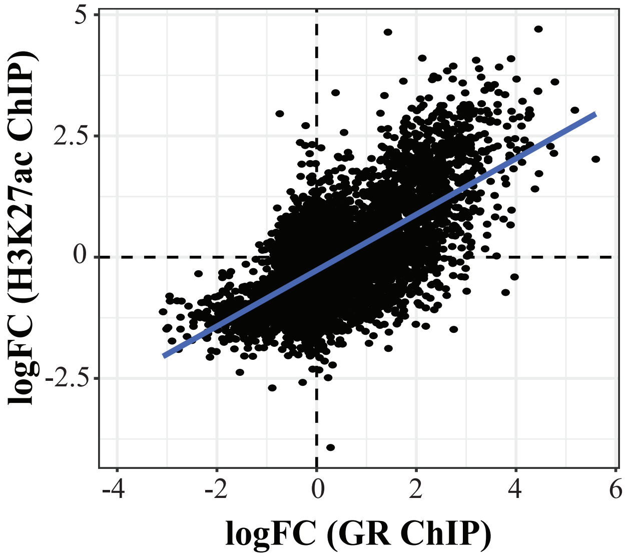

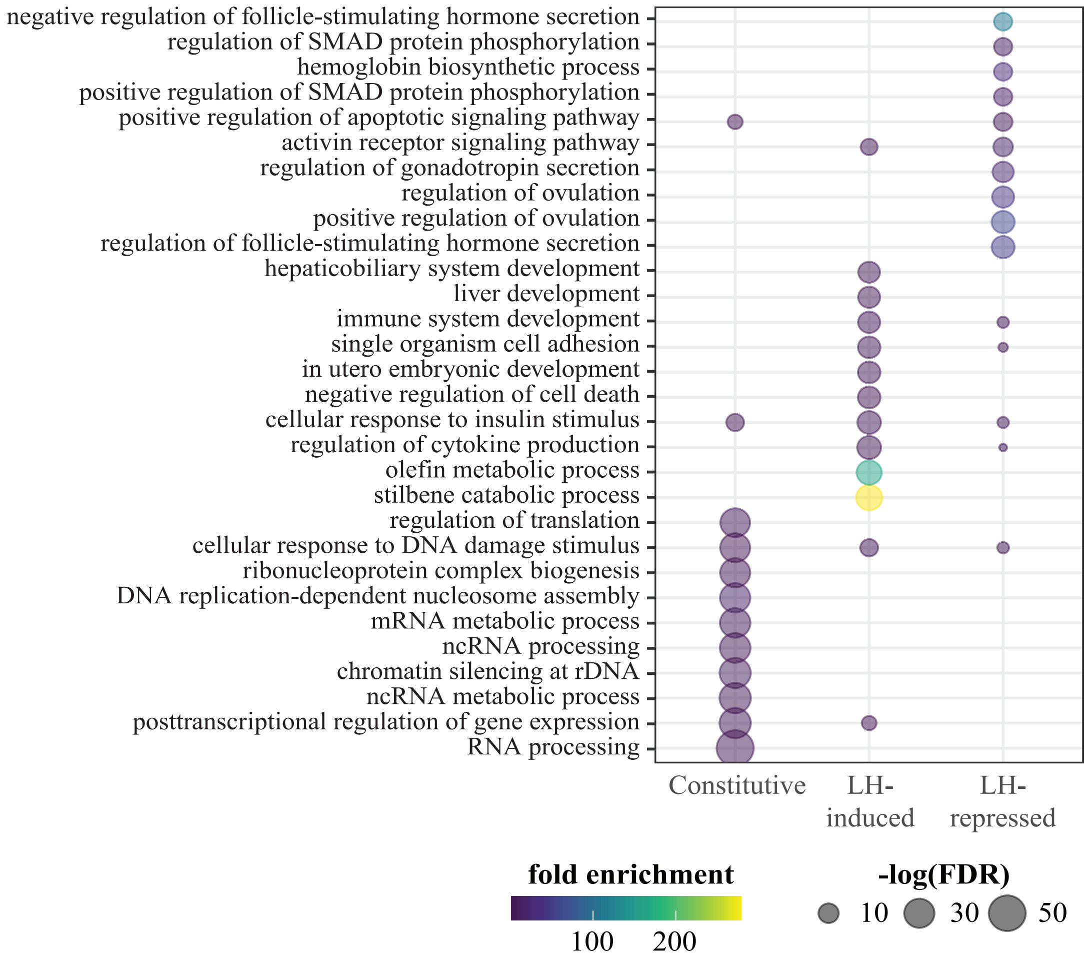

### SFig 5

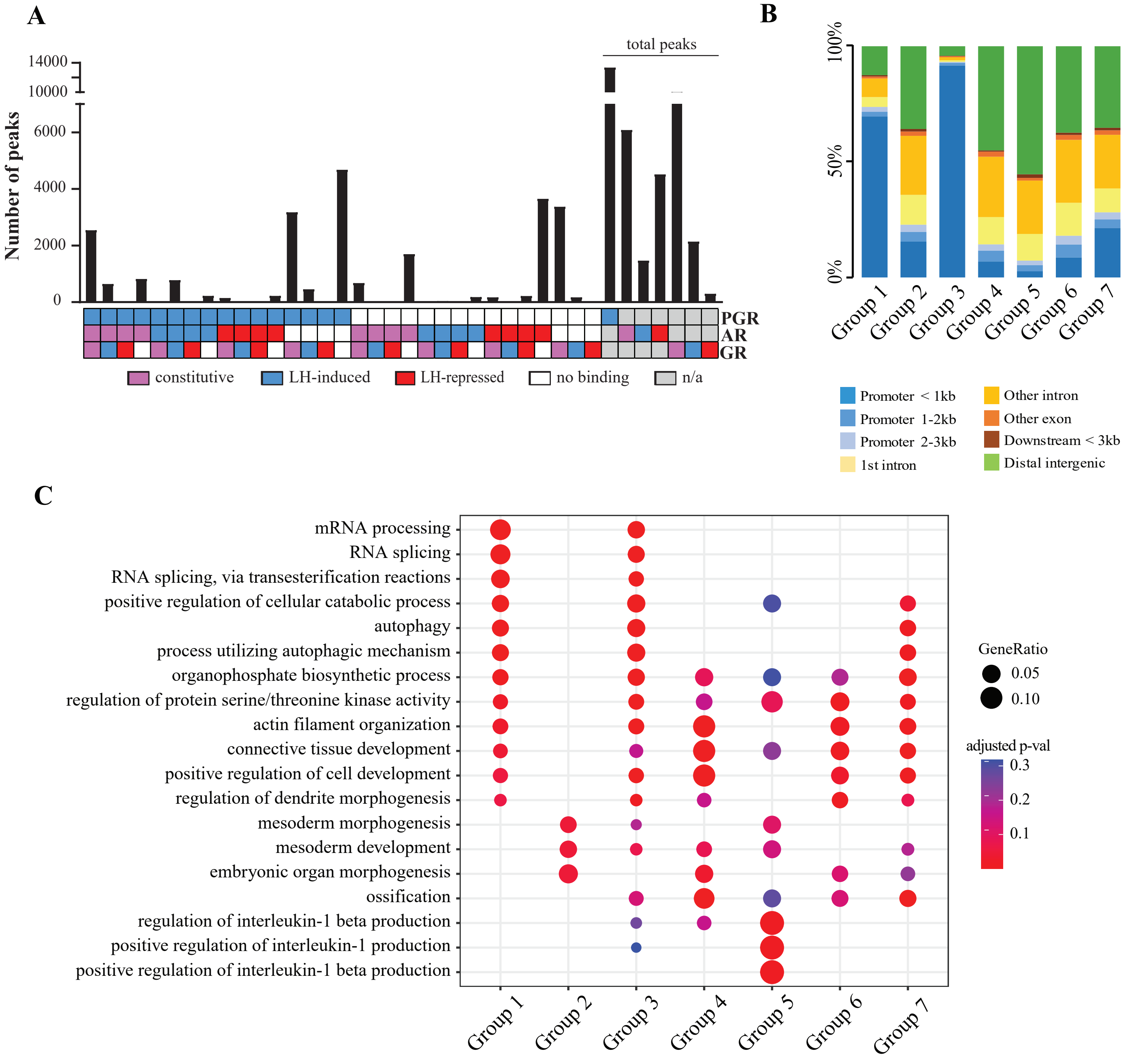

### SFig 6

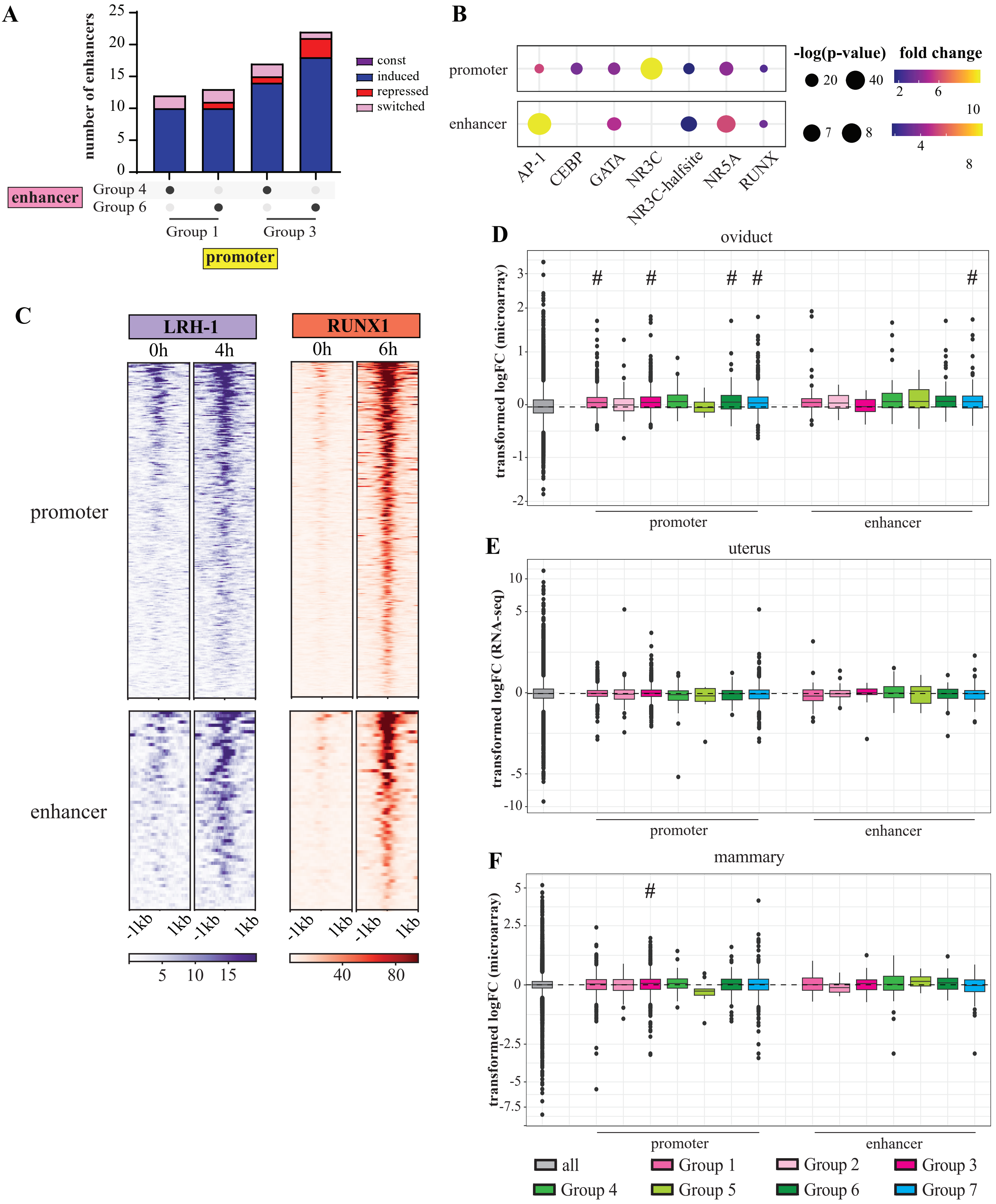
